# Disparate requirements for RAD54L in replication fork reversal

**DOI:** 10.1101/2023.07.26.550704

**Authors:** Mollie E. Uhrig, Neelam Sharma, Petey Maxwell, Platon Selemenakis, Alexander V. Mazin, Claudia Wiese

## Abstract

RAD54L is a DNA motor protein with multiple roles in homologous recombination DNA repair (HR). *In vitro*, RAD54L was shown to also catalyze the reversal and restoration of model replication forks. In cells, however, little is known about how RAD54L may regulate the dynamics of DNA replication. Here, we show that RAD54L restrains the progression of replication forks and functions as a fork remodeler in human cells. Analogous to HLTF, SMARCAL1, and FBH1, and consistent with a role in fork reversal, RAD54L decelerates fork progression in response to replication stress and suppresses the formation of replication-associated ssDNA gaps. Interestingly, loss of RAD54L prevents nascent strand DNA degradation in both BRCA1/2- and 53BP1-deficient cells, suggesting that RAD54L functions in both pathways of RAD51-mediated replication fork reversal. In the HLTF/SMARCAL1 pathway, RAD54L is critical, but its ability to catalyze branch migration is dispensable, indicative of its function downstream of HLTF/SMARCAL1. Conversely, in the FBH1 pathway, branch migration activity of RAD54L is essential, and FBH1 engagement is dependent on its concerted action with RAD54L. Collectively, our results reveal disparate requirements for RAD54L in two distinct RAD51-mediated fork reversal pathways, positing its potential as a future therapeutic target.

## INTRODUCTION

Faithful and complete DNA replication is critical for genome stability (1). Yet, faithful, and complete DNA replication is challenged by genotoxic insult from endogenous and exogenous sources, leading to obstacles in replication fork progression and replication stress. To ameliorate replication stress, cells have acquired several mechanisms that promote the stabilization of stressed replication forks. One such mechanism is through fork reversal (2,3), a process that involves annealing of both nascent and parental DNA strands (4,5). Fork reversal creates a 4-way DNA structure and a double-stranded DNA end, and is associated with a slowdown in fork progression, providing time for lesion bypass and repair (3,6). Efficient fork reversal requires the RAD51 recombinase (2,4,7–9), the key protein in homologous recombination DNA repair (HR), and several ATP-dependent DNA motor proteins of the SWI2/SNF2 family including SMARCAL1, ZRANB3, HLTF, and the F-box DNA Helicase 1 (FBH1) (10–14). Stringent regulation of fork reversal is critical for genome stability and dependent on a growing list of fork reversal and protection factors (2,14). If these factors are dysregulated, fork reversal can lead to unscheduled fork degradation and genome instability (3,14,15). Notably, several key proteins in HR, such as BRCA1 and BRCA2, have critical roles in protecting reversed forks from nucleolytic degradation (3,15,16).

HR is an essential DNA repair pathway that is also required for robust DNA replication (17). HR between sister chromatids ensures that ssDNA gaps are sealed correctly, and that collapsed replication forks are rescued through joint molecule formation with the undamaged sister. Joint molecule formation relies on the DNA strand exchange activity of RAD51 and its auxiliary proteins, including RAD54L (18).

RAD54L is a member of the SWI2/SNF2 family of DNA-dependent ATPases (19) and was shown to have multiple roles in HR (20). During strand invasion, RAD54L utilizes its ATPase activity to convert the synaptic complex into a displacement-loop (21,22). RAD54L also removes RAD51 from heteroduplex DNA to allow access of a DNA polymerase for repair synthesis (23,24). Independent of its ATPase activity, RAD54L functions at the pre-synaptic stage to stabilize the RAD51 filament (23,25–27). *In vitro*, purified RAD54L promotes the branch migration (BM) of Holliday junctions (HJ) and the reversal and restoration of model replication forks (28,29). In cells, however, the role of RAD54L in replication fork dynamics has remained enigmatic.

Previous studies in mouse embryonic stem and HeLa cells have shown that loss of RAD54L does not lead to the nucleolytic degradation of stalled replication forks (15,30,31). The results from these studies have suggested that RAD54L plays no major role as a classical fork protection factor. However, in one of our studies (31) we noticed that RAD54L may play a role in replication fork restraint. As fork restraint is linked to fork reversal (3,32), we wondered if RAD54L may function in fork reversal in human cells.

Here, we show that RAD54L restrains the progression of replication forks in several human cell lines, including cancer and near-normal cells. We show that replication associated ssDNA gaps are significantly more prevalent in the absence of RAD54L, and that treatment of RAD54L-deficient cells with a PARPi exacerbates accelerated replication fork progression, further enhancing the creation of S1 nuclease-sensitive sites. We provide evidence that RAD54L functions in two distinct pathways of RAD51-mediated fork reversal. In the FBH1 pathway, RAD54L’s engagement largely depends on its ability to catalyze BM. In contrast, RAD54L BM activity is dispensable in the HLTF/SMARCAL1 pathway. Collectively, our results identify disparate requirements for RAD54L in two fork reversal pathways and provide new mechanistic insights on the cooperativity between RAD54L and FBH1 in driving fork reversal.

## MATERIAL AND METHODS

### Cell lines, transfections, siRNAs and Western blots

HeLa and MCF7 cells were obtained from ATCC and maintained as recommended. Hs578T cells were a gift from Dr. Joe Gray (OHSU) and maintained as described (33). hTERT RPE-1 cells were provided by Dr. Tingting Yao (CSU) and maintained as described (34). HeLa and Hs578T cells that are knockout (KO) for *RAD51AP1* or *RAD54L*, and HeLa *RAD54L KO* cells ectopically expressing RAD54L were maintained as described previously (27,31,35). DLD1 (*BRCA2* KO) cells and DLD1 cells expressing the wild type BRCA2 protein were maintained as described (36).

The negative control siRNA (Ctrl) and siRNAs targeting RAD54L, BRCA1, or BRCA2 were described earlier (27,37–39) and obtained from IDT (Table S1). For knockdown of HLTF, FBH1, or 53BP1, pools of 3 target-specific siRNAs were purchased from Santa Cruz Biotechnology or IDT (Table S1). SiRNA forward transfections with Lipofectamine RNAiMAX (Thermo Fisher Scientific) were performed on two consecutive days. The concentration of siRNAs in transfections was 20 nM each. Cells were treated with drugs at 96 h after the first transfection.

Western blot analyses were performed according to our standard protocols (40). The following primary antibodies were used: α-RAD51AP1 ((41); 1:6,000), α-RAD54L (F-11; sc-374598; Santa Cruz Biotechnology; 1:500); α-RAD51 (Ab-1; EMD Millipore; 1:3,000), α-PARP1 (ab6079; Abcam; 1:1,000), α-α-Tubulin (DM1A; Santa Cruz Biotechnology; 1:1,000), α-Histone H3 (ab1791; Abcam; 1:10,000), α-HLTF (E9H5I; 45965; Cell Signaling; 1:6,000) α-FBH1 (sc-81563; 1:100), α-BRCA2 (OP95; EMD Millipore; 1:500), α-BRCA1 (MS110; ab16780; Abcam; 1:50), α-PCNA (sc-25280; Santa Cruz Biotechnology; 1:1,000), α-NUCKS1 ((42); 1:10,000), α-MSH2 (ab52266; Abcam; 1:5,000), α-53BP1 (A300-272A; Bethyl Laboratories; 1:10,000). HRP-conjugated goat anti-rabbit or goat anti-mouse IgG (Jackson ImmunoResearch; 1:10,000) were used as secondaries. Western blot signals were acquired using a Chemidoc XRS+ gel imaging system and ImageLab software version 5.2.1 (BioRad).

### Generation of the MCF7 RAD51AP1 and RAD54L KO cells

To generate MCF7 RAD51AP1 KO cells, a cocktail of three different CRISPR/Cas9 knockout plasmids (Santa Cruz Biotechnology (sc-408187)) each encoding Cas9 nuclease and one of three different *RAD51AP1*-specific gRNAs targeting exons 2, 3 or 5/6 (Table S2) was used to transfect parental MCF7 cells. To generate MCF7 RAD54L KO cells, a combination of two *RAD54L* CRISPR/Cas9-nic KO plasmids each containing one of two different sgRNAs (*i.e*., sgRNA (54L)-A and sgRNA (54L)-B; Santa Cruz Biotechnology (sc-401750-NIC); Table S2) was used. Disruption of *RAD51AP1* or *RAD54L* was validated by sequence analysis after genomic DNA was isolated from a selection of edited and non-edited clonal isolates using DNeasy Blood & Tissue Kit (Qiagen). *RAD51AP1* and *RAD54L* genomic DNA sequences were amplified by PCR primers flanking the sgRNA target sites (Table S3). PCR products were gel purified, cloned into pCR4-TOPO (Invitrogen) and transformed into TOP10 competent *E. coli*. Plasmid DNA was prepared using ZR Plasmid Miniprep-Classic Kit (Zymo Research) and submitted for sequencing. For each cell line, 15-20 individually cloned amplicons were analyzed by Sanger sequencing.

### iPOND assay

The iPOND assay was performed as described (43). Briefly, 1.5×10^6^ HeLa cells were plated in eight 145 mm plates 72 h prior to treatment. Cells were pulse-labeled in 10 µM EdU (Invitrogen) for 20 min and washed twice in PBS. Cells were either fixed immediately or incubated for 2 h in medium containing 3 mM HU. Cells were fixed in 10 ml 1% paraformaldehyde for 10 min prior to quenching the reaction by addition of 1 ml 1.25 M glycine. The cells were washed three times in PBS and harvested using a cell scraper. Cells were spun at 1000×g and 4ᵒC, washed in ice-cold PBS twice, and then flash frozen. To begin protein extraction, cells were permeabilized in 1.5 ml 0.25% Triton X-100/PBS at RT for 30 min. Cell were washed once in 0.5% BSA/PBS and once in PBS before performing the Click-iT reaction with Biotin-Azide (Thermo Fisher Scientific) at RT for 2 h. Then, the cells were pelleted, washed once in 0.5% BSA/PBS and once in PBS, and lysed in lysis buffer (50 mM Tris-HCl [pH 8.0], 1% SDS, and protease/phosphatase inhibitors (Thermo Fisher Scientific)). Cells were sonicated at high setting with 30 s on/off cycles using a Bioruptor UCD-200 (Diagenode). Lysates were cleared by centrifugation at 14000×g. Cleared lysates were diluted with PBS containing protease and phosphatase inhibitors to reduce the final SDS concentration to 0.5% and then incubated with 80 µl equilibrated streptavidin agarose beads (Novex) at 4ᵒC for 16 h. Beads were washed twice with chilled lysis buffer and once with 1M NaCl/PBS before bound proteins were eluted in 80 µl 2×LDS and 95ᵒC for 45 min. Captured proteins were fractionated on NuPAGE gels (Thermo Fisher Scientific) for immunoblot analyses.

### Site-directed mutagenesis and lentiviral transduction

Mutations in RAD54L were generated in pENTR1A-RAD54L-HA (31) using the Q5 Site-Directed Mutagenesis Kit (New England Biolabs) (Table S4). pENTR1A constructs were transferred into pLentiCMV/TO DEST#2 (44) using Gateway LR Clonase II (Thermo Fisher Scientific) for the production of lentiviral particles in HEK293FT cells (Thermo Fisher Scientific), as described (44). Lentivirus was used to transduce HeLa *RAD54L* KO cells in 6 µg/ml polybrene, as described (31,44).

### DNA fiber assay

The DNA fiber assay was performed as described (31). In the modified fiber assay with S1 nuclease, cells were labeled for 25 min each with 25 µM CldU and 250 µM IdU followed by 10 min in 200 µM thymidine, as described (45). Cells were harvested in cold PBS, mixed with unlabeled cells (1:1), and centrifuged at 1,500×g for 5 min at 4°C. Cells were resuspended in hypotonic buffer (10 mM HEPES [pH 7.5], 150 mM NaCl, 0.3 M sucrose, and 0.5% Triton-X-100) and incubated on ice for 15 min before centrifugation as above. Cells were resuspended in S1-nuclease buffer (30 mM sodium acetate [pH 4.6], 10 mM zinc acetate, 5% glycerol, and 50 mM NaCl) with or without 10U/mL S1 nuclease enzyme (Thermo Fisher Scientific) and incubated for 10 min at 37°C. Cells were centrifuged as above and resuspended in 50 µl lysis buffer (0.5% SDS, 200 mM Tris pH 7.5, 50 mM EDTA). The fibers were spread, imaged, and measured as described (10). Two slides per sample were prepared for each experimental repeat, and each pair of slides was blinded after immunodetection to avoid bias.

Fiber tract lengths in HeLa cell lines and hTERT RPE-1 cells were measured to evaluate the consequences of mild replication stress, as described (32). To do so, cells were pulse labeled with CldU for 20 min followed by IdU containing 25 µM hydroxyurea (Sigma) or 25 nM camptothecin (Sigma) for 20 min (HeLa cells), or 30 min (Hela and hTERT RPE-1 cells). Only IdU tracts following a CldU tract were analyzed.

### Comet assay

The Comet assay (Trevigen) was performed as recommended by the manufacturer. Cells seeded in 60-mm plates were incubated for 48 hours before 5-hour treatment with 4 mM HU. Cells were harvested in cold PBS and combined with molten low melting (LM) agarose at 1:10. Fifty µL of this mixture were spread on comet slides and incubated for 30 min at 4°C in the dark. The slides were then incubated overnight at 4°C in lysis buffer. Following a 30 min incubation in neutral comet electrophoresis buffer (10 mM Tris base, 250 mM sodium acetate) at 4°C, electrophoresis was performed at 25V and 4°C for 30 min in neutral electrophoresis buffer. The slides were then incubated in DNA precipitation solution (7.5 M NH_4_Ac and 95% ethanol) for 30 min followed by 30 min incubation in 70% ethanol. Slides were dried at 37°C for 1 hour and stained with 0.3× SYBR Gold (Thermo Fisher). Images were acquired on a Zeiss Axio-Imager.Z2 microscope equipped with Zen Blue software (Carl Zeiss Microscopy) using a 20× objective, and 100 comets were measured per condition. The lengths of the comet tails were measured using ImageJ software (https://imagej.net).

### Statistics and reproducibility

GraphPad Prism 9 software was used to perform statistical analyses on data obtained from 3 independent experiments. To assess statistical significance ANOVA tests were performed as indicated, and *P* ≤ 0.05 was considered significant.

## RESULTS

### RAD54L is recruited to stalled DNA replication forks

To begin exploring the involvement of RAD54L in replication fork progression in human cells, we applied the isolation of proteins on nascent DNA (iPOND) assay (43). We show that RAD54L is enriched at replication forks upon a 2 h-treatment of cells with 3 mM hydroxyurea (HU) (Fig. 1A, B), in accord with results from others (30). Enrichment of RAD54L mirrored that of its binding partners RAD51 and NUCKS1. In contrast, enrichment of PCNA was downregulated after HU, as expected. These results demonstrate that RAD54L is recruited to stalled replication forks.

**Fig. 1.**
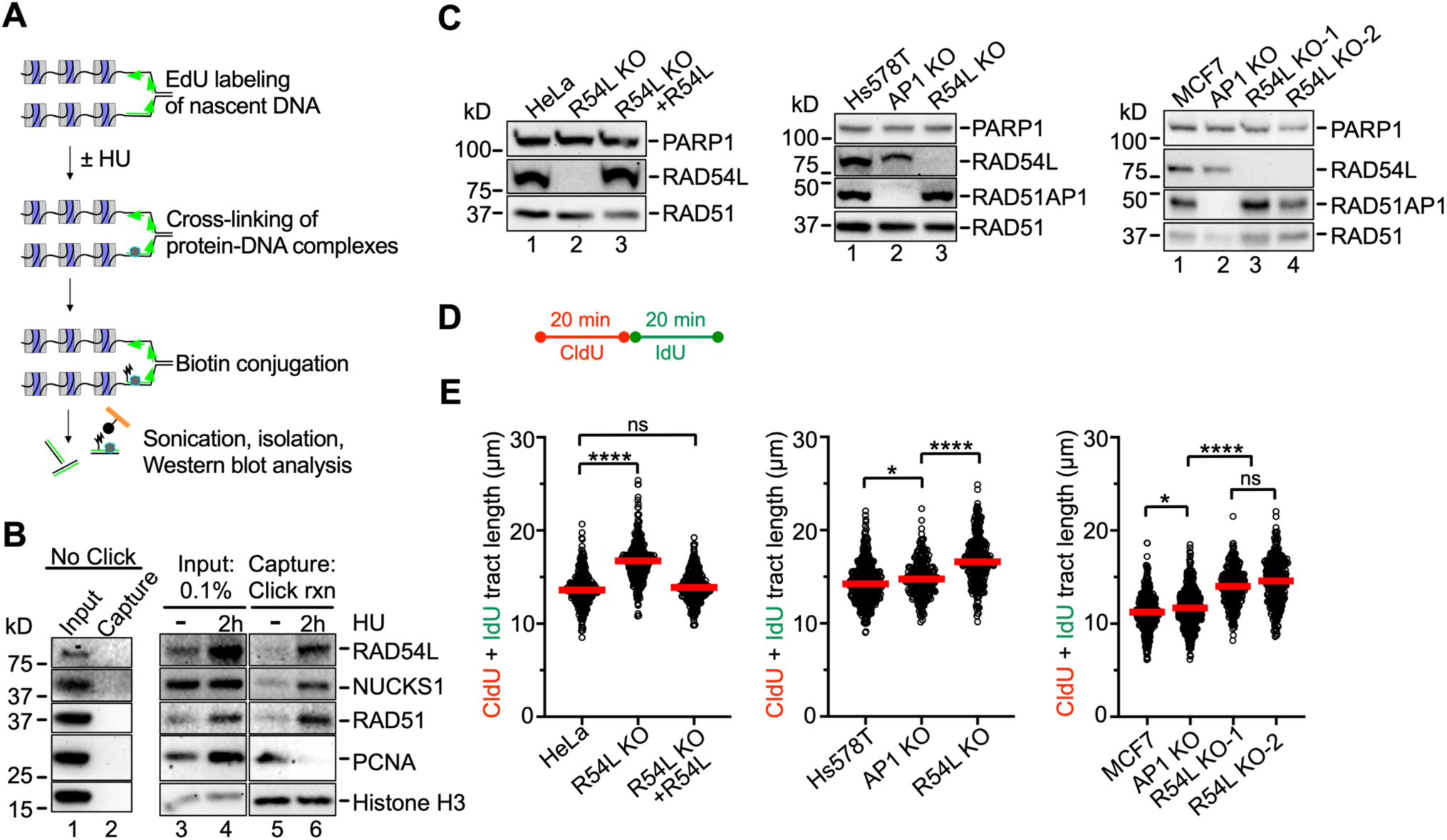
RAD54L is enriched at stalled replication forks and restrains fork progression. (**A**) Schematic illustration of the iPOND assay. (**B**) Western blots of the input and iPOND samples probed for proteins, as indicated. NUCKS1 is a RAD54L interacting protein (27). Click rxn, Click reaction; the “no Click” condition represents cells pulsed with EdU and processed without biotin-azide in the Click reaction step. (**C**) Western blots of HeLa, R54L KO, and R54L KO cells ectopically expressing R54L-HA. (**D**) Schematic of the protocol for the DNA fiber assay to determine replication tract lengths. (**E**) Dot plot with medians of CldU+IdU tract lengths in parental cells and R54L KO derivatives generated in HeLa, Hs578T, and MCF7 cells. RAD51AP1 KOs (AP1) and the R54L KO ectopically expressing R54L-HA (31) are shown for comparison purposes (n=3; at least 100 fiber tracts/experiment analyzed). Fiber tract lengths were analyzed by Kruskal-Wallis test followed by Dunn’s multiple comparisons test (ns, not significant; *, p<0.05; ****, p<0.0001). R54L: RAD54L.

### RAD54L restrains replication fork progression in unperturbed cells

Previously, we generated HeLa RAD54L KO cells and RAD54L KO cells ectopically expressing HA-tagged RAD54L, which fully rescues the sensitivity of RAD54L KO cells to mitomycin C (MMC) and Olaparib (31). Using the single molecule DNA fiber assay, we noticed that RAD54L KO cells had longer replication tracts than the controls (31). These results were independently replicated here (Fig. 1C-E; Fig. S1A). We also previously generated RAD54L KOs in Hs578T cells (31) and in this study in MCF7 cells (Fig. 1C; Table S5). Both Hs578T RAD54L KO and MCF7 RAD54L KO cells have longer replication tracts than parental cells (Fig. 1E; Fig. S1B, C). Moreover, as in the HeLa cell derivatives (31), fiber tracts in the RAD54L KOs were significantly longer than those in Hs578T or MCF7 cells deficient in RAD51AP1 (Fig.1E; Fig. S1B, C). RAD51AP1, like RAD54L, is also a RAD51-associated HR protein that enhances the activity of the RAD51 recombinase (46,47). These results show that unperturbed DNA replication proceeds faster in the absence of RAD54L and that this effect is not cell-type specific.

### RAD54L is dispensable for the recovery of cells from stalled DNA replication

To understand the fate of stalled replication forks, we treated the Hs578T cell derivatives with 4 mM HU for 5 h, which blocks DNA synthesis and stalls replication fork movement (14). Using the DNA fiber assay, we then monitored the recovery of cells from stalled replication (Fig. 2A). We determined the ability of Hs578T cells and derivatives to restart DNA replication by measuring the lengths of IdU tracts proceeded by a CldU tract. The results show that RAD54L-deficient Hs578T cells recover as fast as their parental cells from stalled DNA replication (Fig. 2B, right panel; Fig. S2A). In contrast, RAD51AP1-deficient Hs578T cells are significantly impaired in fork restart. These results mirror our published results generated in HeLa cells (31) and demonstrate that RAD54L is dispensable for the recovery from stalled DNA replication in human cells.

**Fig. 2.**
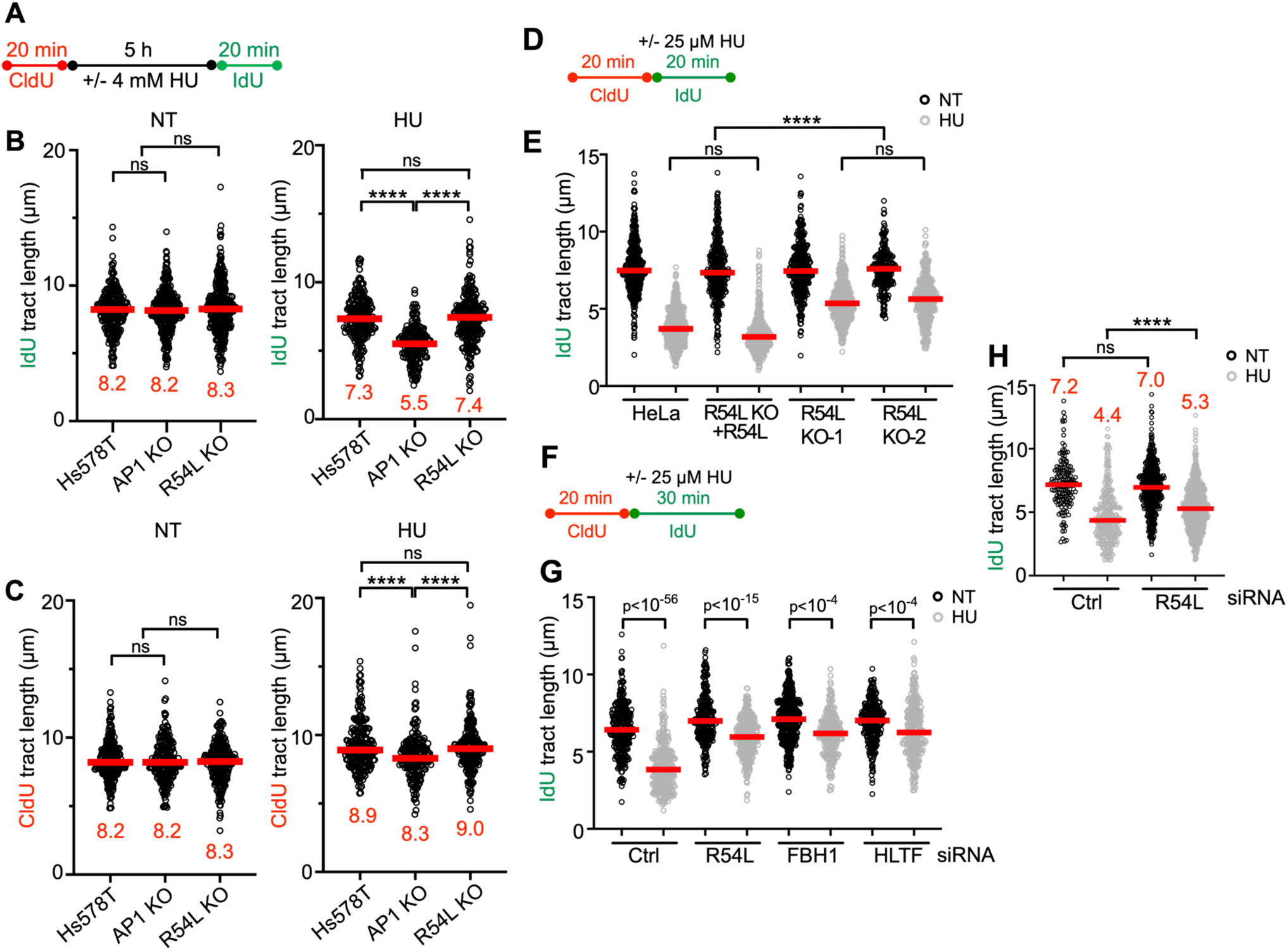
Loss of RAD54L does not affect replication restart and leads to longer replication tracts during mild replication stress. (**A**) Schematic of the DNA fiber assay protocol used to assess replication restart. (**B**) Dot plot with medians of IdU tract lengths in untreated (NT) or HU treated Hs578T and AP1 or R54L KO cells. (**C**) Dot plot with medians of CldU tract lengths in untreated (NT) or HU treated Hs578T and AP1 or R54L KO cells. (n=3; at least 100 fiber tracts/experiment analyzed). (**D**) Schematic of the DNA fiber assay protocol used to evaluate the progression of replication forks during mild replication stress in E. (**E**) Dot plot with medians of IdU tract lengths in HeLa control cells, R54L KO+R54L, and two independently isolated R54L KO cell lines treated with or without 25 μM HU during the IdU pulse (n=3; at least 100 fiber tracts/experiment analyzed). (**F**) Schematic of the DNA fiber assay protocol used to evaluate the progression of replication forks during mild replication stress in G, H. (**G**) Dot plot with medians of IdU tract lengths in HeLa cells transfected with control (Ctrl), R54L, FBH1, or HLTF siRNA and treated with or without 25 μM HU during the IdU pulse (n=3; at least 100 fiber tracts/experiment analyzed). (**H**) Dot plot with medians of IdU tract lengths in hTERT RPE-1 cells transfected with Ctrl or R54L siRNA and treated with or without 25 μM HU during the IdU pulse (n=3; at least 100 fiber tracts/experiment analyzed). Fiber tract lengths were analyzed by Kruskal-Wallis test followed by Dunn’s multiple comparisons test (ns, not significant; ****, p<0.0001, or as indicated). AP1: RAD51AP1; R54L: RAD54L.

### RAD54L slows fork progression upon nucleotide depletion

In response to replication stress, replication forks reverse into four-way junctions through annealing of the nascent DNA strands (2,9). Reversed forks must be protected from nucleolytic attack to prevent fork attrition (10,13,15,48,49). To assess if RAD54L in Hs578T cells functions in the protection of replication forks from unprogrammed nuclease degradation, CldU tracts in cells exposed to HU were measured. The results show that CldU tracts in Hs578T RAD54L KO cells are not shorter than in parental cells (Fig. 2C, right panel; Fig. S2A). Hence, RAD54L appears to play no major role in protecting reversed forks from nuclease attrition in cells with otherwise unaltered fork protection pathways. These results are in accord with earlier studies by us and others using different cell types (15,30,31).

Previous studies have shown that a defect in replication fork restraint is linked to compromised fork reversal (2,32,50,51). It is also known that RAD54L has branch migration (BM) activity and can reverse model replication forks *in vitro* (28,29,52). Prompted by our findings that identified a defect in replication fork restraint in RAD54L-deficient cells (Fig. 1E), we monitored the progression of replication forks in HeLa cell derivatives under a low concentration of HU (25 µM) given within the IdU pulse (Fig. 2D). In these experiments, we measured the lengths of IdU tracts following a CldU tract. The results show that the replication tracts in two independently isolated RAD54L KO lines (31) are significantly longer than those in parental cells or RAD54L KO cells ectopically expressing RAD54L (Fig. 2E; Fig. S2B). These results strongly suggest that in response to HU, which induces fork reversal (48), accelerated fork progression is a consequence of RAD54L deficiency. These results also suggest that RAD54L may catalyze the reversal of replication forks *in vivo*.

Next, we used a similar protocol of the DNA fiber assay with extended IdU tract length (Fig. 2F) to compare the consequences of RAD54L deficiency to loss of HLTF or FBH1, two established fork remodelers that each function in a different RAD51-mediated fork reversal pathway (14,32,51,53,54). To do so, we depleted RAD54L, HLTF, or FBH1 in HeLa cells (Fig. S2C) and measured IdU tract lengths in the presence of 25 µM HU. As observed previously in different cell types (14,32), loss of HLTF or FBH1 expression led to significantly longer replication tracts than those in control cells (Fig. 2G; Fig. S2D). Of note, after depletion of RAD54L, IdU tracts were also significantly longer than in control cells (Fig. 2G; Fig. S2D).

Last, we tested if loss of RAD54L expression would lead to a defect in fork restraint in non-cancerous hTERT RPE-1 cells. To do so, we depleted RAD54L in hTERT RPE-1 cells (Fig. S2E) and measured IdU tract lengths with the fiber assay (Fig. 2H). Compared to control cells transfected with a non-depleting Ctrl siRNA, IdU tracts were significantly longer in RAD54L-deficient hTERT RPE-1 cells (Fig. 2H; Fig. S2F). Collectively, our results show that RAD54L’s ability to restrain fork progression is not confined to cancer cell lines and occurs in both unperturbed cells (Fig. 1E) and upon exposure of cells to mild replication stress (Fig. 2E, G, H).

### Accelerated fork progression in RAD54L-deficient cells is associated with ssDNA gap formation

Treatment of cells with a PARP1 inhibitor (PARPi) leads to accelerated replication fork progression, impediments in nascent DNA strand maturation, and fork collapse (55–58). RAD54L-deficient cells are sensitive to the PARPi Olaparib (31,34), and treatment of RAD54L KO cells with Olaparib further exacerbates fork progression (Fig. 3A, B; Fig. S3A; (31)). Expression of ectopic RAD54L in RAD54L KO cells reverses accelerated fork progression, as expected (Fig. 3B; Fig. S3A).

**Fig. 3.**
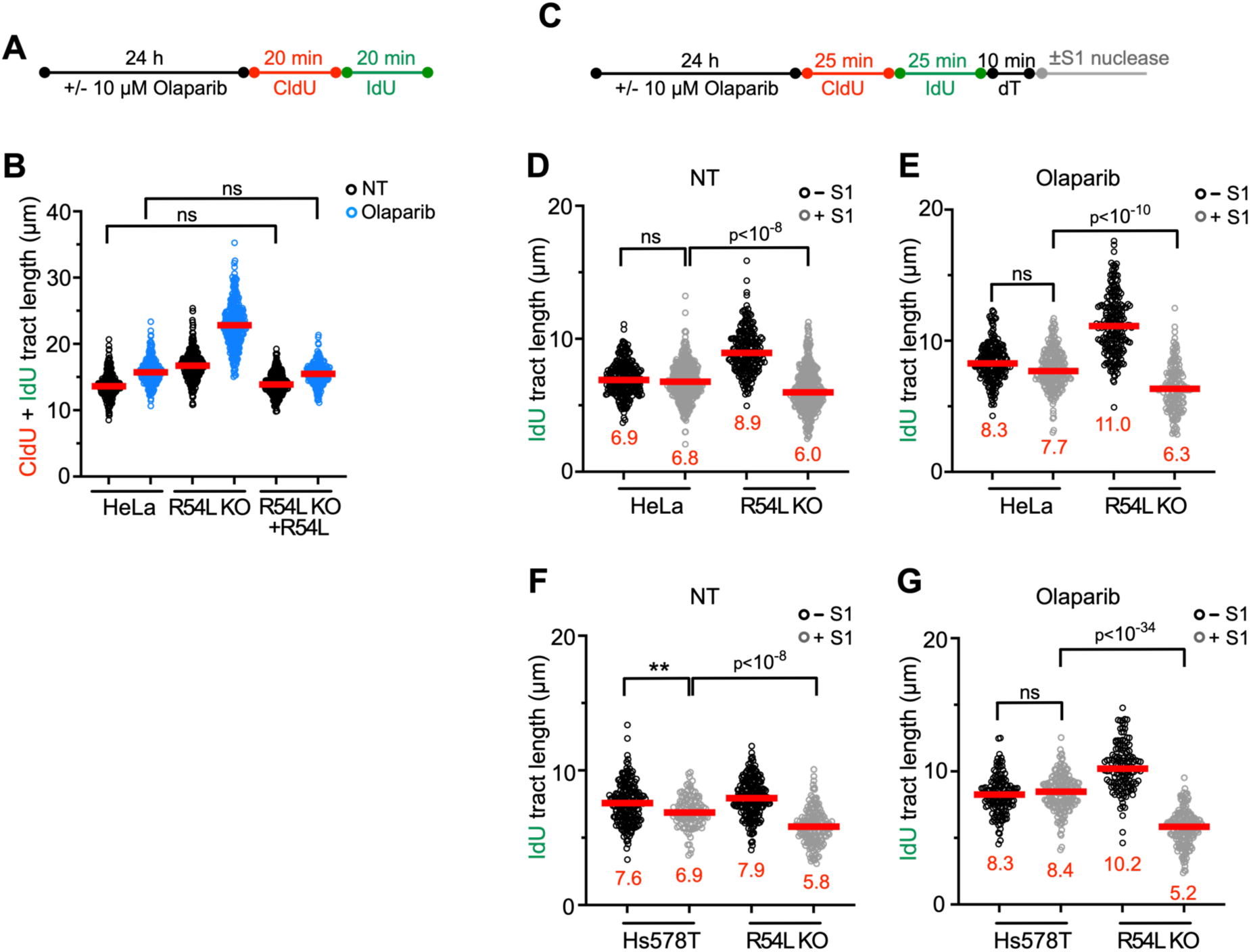
Loss of RAD54L accelerates replication fork progression and ssDNA gap formation. (**A**) Schematic of the DNA fiber assay protocol to assess fork progression with and without Olaparib. (**B**) Dot plot with medians of CldU+IdU fiber tract lengths in HeLa, R54L KO, and R54L KO+R54L cells with or without Olaparib (n=3; at least 100 fiber tracts/experiment analyzed). (**C**) Schematic of the modified DNA fiber assay protocol with S1 nuclease in unperturbed and Olaparib treated cells. (**D**) Dot plot with medians of IdU fiber tract lengths in untreated (NT) HeLa and R54L KO cells with or without S1 nuclease (n=3; at least 100 fiber tracts/experiment analyzed). (**E**) Dot plot with medians of IdU fiber tract lengths in HeLa and R54L KO cells treated with Olaparib and with or without S1 nuclease (n=3; at least 100 fiber tracts/experiment analyzed). (**F**) Dot plot with medians of IdU fiber tract lengths in untreated (NT) Hs578T and R54L KO cells with or without S1 nuclease (n=3; at least 100 fiber tracts/experiment analyzed). (**G**) Dot plot with medians of IdU fiber tract lengths in Hs578T and R54L KO cells treated with Olaparib and with or without S1 nuclease (n=3; at least 100 fiber tracts/experiment analyzed). Fiber tract lengths were analyzed by Kruskal-Wallis test followed by Dunn’s multiple comparisons test (ns, not significant; (**, p<0.01; ****, p<0.0001). R54L: RAD54L.

To test for the presence of ssDNA formed at replication gaps, we used S1 nuclease, which cleaves replication intermediates that contain ssDNA (59). We incubated control cells and Olaparib treated cells with and without S1 nuclease (Fig. 3C). In these experiments, we assessed the derivatives of both HeLa and Hs578T cells and found that S1 treatment resulted in more prominently shortened replication tracts in cells deficient in RAD54L (Fig. 3D, F; Fig. S3A, B). Treatment of cells with Olaparib and S1 nuclease exacerbated replication tract shortening in RAD54L KO cells (Fig. 3E, G; Fig. S3A, B). These results show that replication in the absence of RAD54L is not only accelerated but also proceeding with elevated production of ssDNA gaps. Moreover, treatment of RAD54L KO cells will Olaparib further increases the abundance of ssDNA gaps.

### Loss of RAD54L restores fork stability in both BRCA1/2- and 53BP1-deficient cells

The fork reversal activities of SMARCAL1, ZRANB3, and HTLF lead to the degradation of nascent strand DNA in BRCA1/2-deficient cells upon treatment with HU (9,10). Given our results that suggest that RAD54L may function similarly to these established fork remodelers, we tested if RAD54L loss prevents nascent strand DNA degradation in BRCA1/2-deficient cells (Fig. 4A). We depleted BRCA2 in HeLa, RAD54L KO, and RAD54L-rescued cell lines (Fig. 4B) and measured fork degradation by IdU/CldU ratios in response to a 5 h treatment of cells with 4 mM HU. As expected, BRCA2 knockdown in HeLa and RAD54L KO+RAD54L cells led to significantly reduced IdU/CldU tract ratios (p<0.0001; Fig. 4C; Fig. S4A). In contrast, forks in RAD54L KO cells with BRCA2 knockdown withstood fork attrition (Fig. 4C; Fig. S4A). Similarly, IdU/CldU tract ratios in HeLa cells with BRCA1 knockdown were reduced, while no such effect was observed in RAD54L KO cells with BRCA1 knockdown (Fig. S4B-D). These results show that RAD54L loss in BRCA1/2-deficient HeLa cells prevents nascent strand DNA degradation in response to HU.

**Fig. 4.**
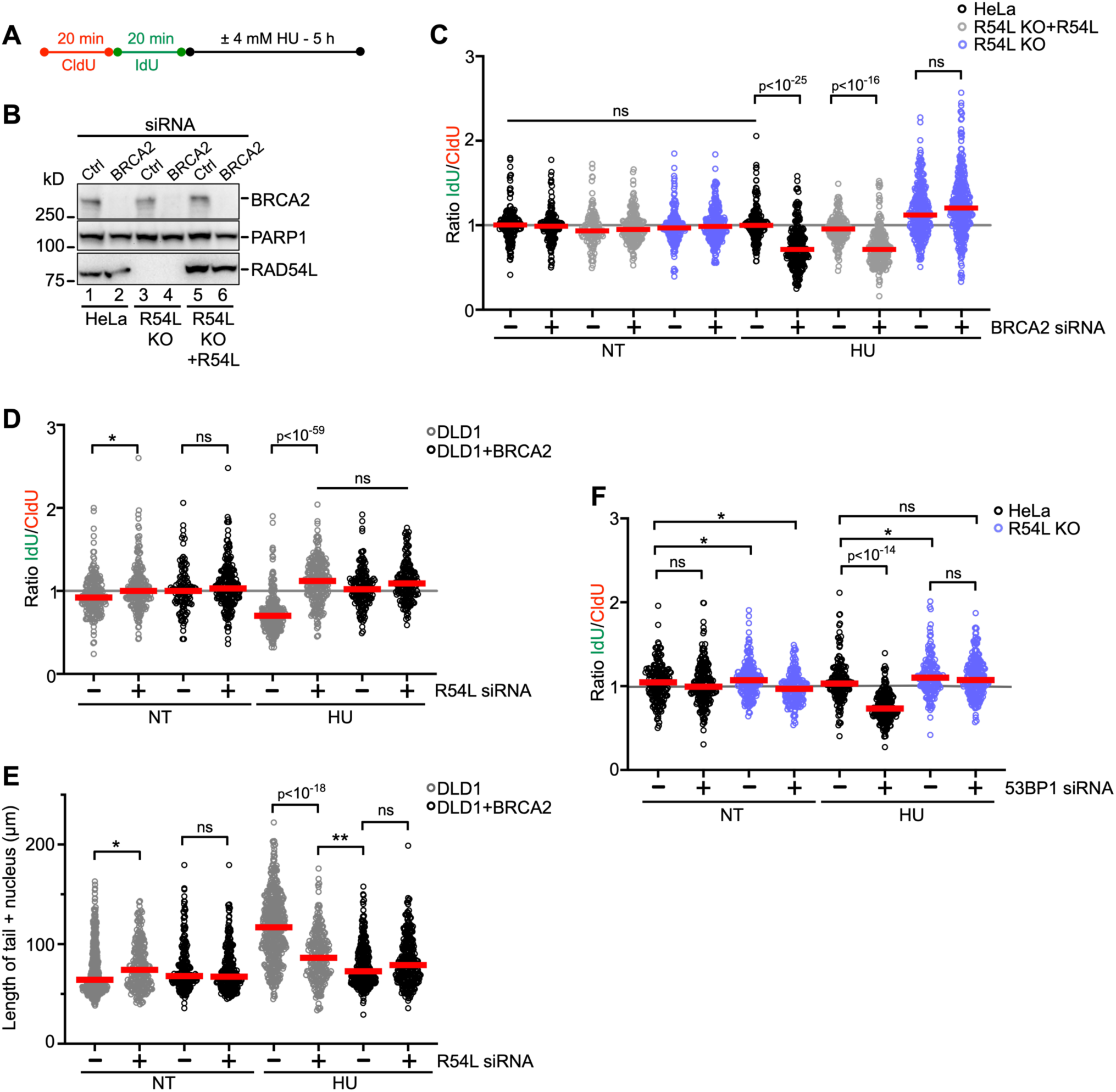
Loss of RAD54L prevents fork degradation in both BRCA2- and 53BP1-deficient cells. (**A**) Schematic of the DNA fiber assay protocol to assess fork degradation. (**B**) Representative Western blots to show extent of BRCA2 knockdown in HeLa cells and derivatives for the experiments shown in C. Loading control: PARP1. (**C**) Dot plot with medians of IdU/CldU tract length ratios in HeLa, R54L KO, and R54L KO+R54L cells transfected with Ctrl (-) or BRCA2 siRNA and treated with or without HU (n=3; at least 100 fiber tracts/experiment analyzed). (**D**) Dot plot with medians of IdU/CldU tract length ratios in DLD1 and DLD1+BRCA2 cells transfected with Ctrl (**-**) or RAD54L siRNA and treated with or without HU (n=3; at least 100 fiber tracts/experiment analyzed). (**E**) Dot plot with medians after neutral Comet assay in DLD1 and DLD1+BRCA2 cells transfected with Ctrl (**-**) or RAD54L siRNA and treated with or without HU (n=3; at least 100 Comet tails/experiment analyzed). (**F**) Dot plot with medians of IdU/CldU tract length ratios in HeLa and R54L KO cells transfected with Ctrl (**-**) or 53BP1 siRNA and treated with or without HU (n=3; at least 100 fiber tracts/experiment analyzed). IdU/CldU ratios and lengths of Comet signals were analyzed by Kruskal-Wallis test followed by Dunn’s multiple comparisons test (ns, not significant; *, p<0.05; **, p<0.01, or as indicated). R54L:RAD54L. NT: not treated.

Next, we used DLD1 (BRCA2 KO) cells and a DLD1 cell line stably expressing the wild type BRCA2 protein (36) and depleted RAD54L in these two cell lines (Fig. S4E). IdU/CldU tract length ratios were significantly reduced in DLD1 cells after treatment with HU (Fig. 4D; Fig. S4F), as shown previously (36). In contrast, knockdown of RAD54L in HU-treated DLD1 cells gave rise to IdU/CldU tract ratios with a distribution not significantly different to that from DLD1+BRCA2 cells (Fig. 4D; Fig. S4F). These results show that RAD54L loss in BRCA2-deficient DLD1 cells prevents nascent strand DNA degradation after treatment of cells with HU.

To assess the impact of RAD54L deficiency on DSB formation in DLD1 cells, we subjected DLD1 and DLD1+BRCA2 cells (transfected with Ctrl or RAD54L siRNA) to HU and DSB detection by neutral comet assay. Consistent with blocked fork degradation (Fig. 4D), depletion of RAD54L in DLD1+BRCA2 cells had no significant impact on DSB formation (Fig. 4E; Fig. S4G). In contrast, depletion of RAD54L in DLD1 (BRCA2 KO) cells led to significantly reduced DSB formation (Fig. 4E; Fig. S4G). These results show that RAD54L activity contributes to the formation of DSBs in BRCA2-deficient DLD1 cells treated with HU.

Aside from the SMARCAL1/HLTF/ZRANB3 pathway, a second RAD51-dependent fork reversal pathway has been described. This pathway relies on the FBH1 DNA helicase, and the 53BP1 protein is one critical fork protection factor in this pathway (14,54). Using HeLa and RAD54L KO cells, we tested if RAD54L activity would also contribute to fork degradation in 53BP1-depleted cells (Fig. S4H). Upon treatment of 53BP1-depleted HeLa cells with HU, IdU/CldU tract length ratios were reduced (p<0.0001; Fig. 4F; Fig. S4I), indicative of fork attrition. However, IdU/CldU ratios in HU-treated 53BP1-depleted RAD54L KO cells were not different from those in RAD54L KO cells transfected with Ctrl siRNA (Fig. 4F; Fig. S4I). Collectively, our results show that RAD54L activity contributes to nascent strand DNA degradation in both BRCA1/2- and 53BP1-deficient cells, suggesting that RAD54L functions in each of the two described RAD51-mediated fork reversal pathways.

### FBH1 activity in fork reversal is dependent on RAD54L

To better understand how RAD54L is engaged in each of the two RAD51-mediated fork reversal pathways, we depleted FBH1, HLTF, or SMARCAL1 in both HeLa and RAD54L KO cells (Fig. S5A) and monitored the lengths of IdU tracts spontaneously and under mild, HU-induced replication stress (Fig. 5A). In untreated cells, IdU tract lengths in HeLa and RAD54L KO cells transfected with Ctrl, FBH1, or HLTF siRNA were not significantly different from each other (p>0.999; Fig. S5B, C). SMARCAL1 knockdown led to longer IdU tracts in untreated RAD54L KO cells than in parental cells (p<0.0001; Fig. S5B, C). As expected and indicative of ongoing fork reversal, IdU tract lengths were significantly shorter in HU-treated HeLa cells than in untreated HeLa cells (p<0.0001; Fig. 5B; Fig. S5C). After HU, loss of either HLTF or SMARCAL1 led to longer IdU fiber tracts in both HeLa and RAD54L KO cells due to compromised fork restraint. Compared to the equivalent knockdown condition in HeLa cells, loss of either HLTF or SMARCAL1 in RAD54L KO cells affected IdU fiber tract lengths only moderately (HLTF; p=0.043) or not at all (SMARCAL1; p>0.999), suggesting that both HLTF and SMARCAL1 may function independently or upstream of RAD54L. In contrast, IdU tract lengths in RAD54L KO cells with FBH1 knockdown were significantly shorter than those in HeLa cells with FBH1 knockdown (p<0.0001; Fig. 5B), suggesting that the activity of FBH1 in fork reversal is dependent on RAD54L.

**Fig. 5.**
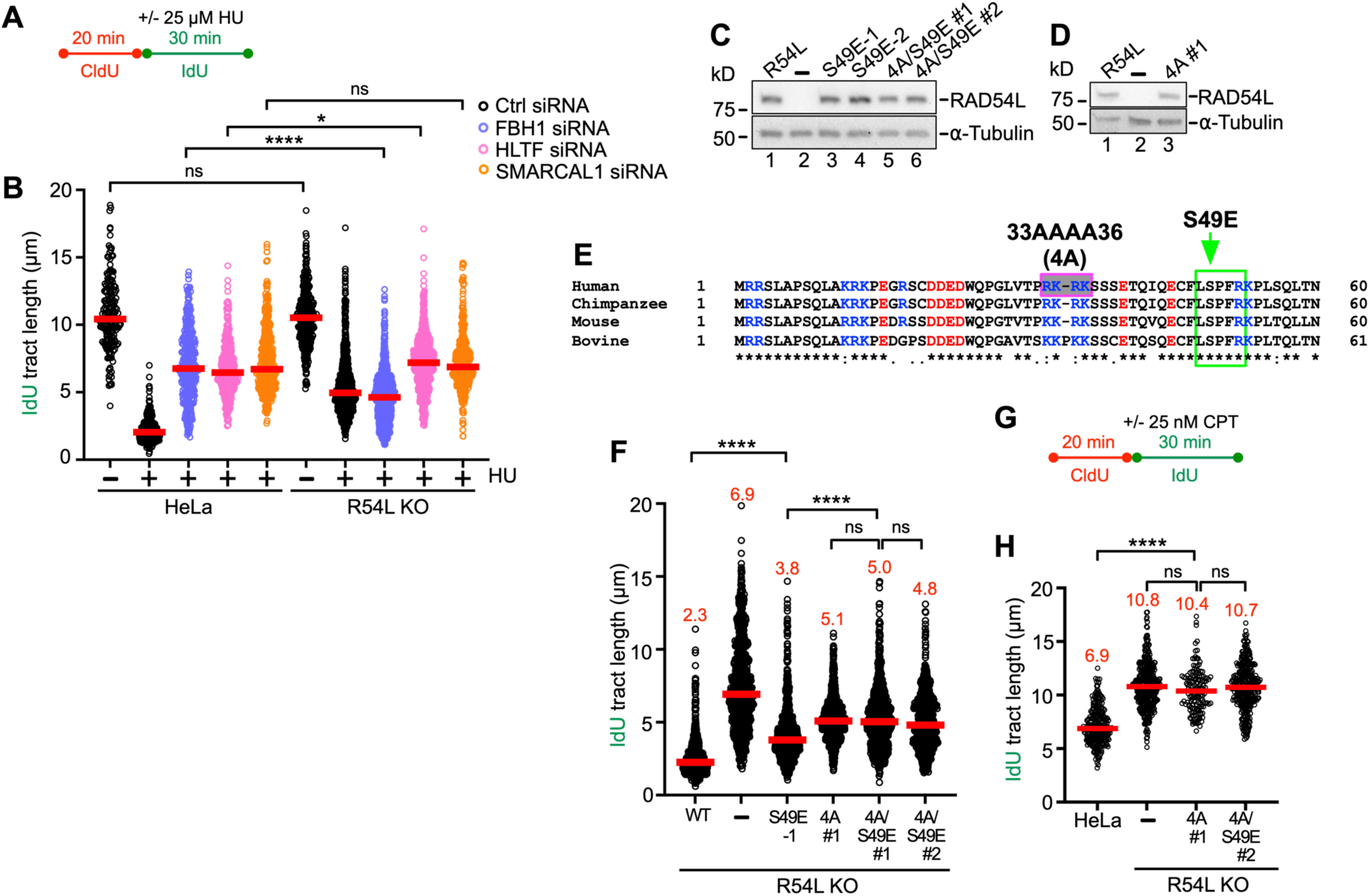
RAD54L’s branch migration activity contributes to its ability to restrain fork progression. (**A**) Schematic of the DNA fiber assay protocol used in B, F. (**B**) Dot plot with medians of IdU tract lengths in HeLa and R54L KO cells transfected with Ctrl, FBH1, HLTF, or SMARCAL1 siRNA (n=3; at least 100 fiber tracts/experiment analyzed). (**C**) Representative Western blots to show expression of wild type R54L (lane 1) and R54L mutant protein (lanes 3-6) in HeLa R54L KO cells. R54L-S49E expressing lines are clonal isolates; R54L-4A/S49E expressing cells are puromycin-resistant cell populations. Loading control: ⍺-Tubulin. (**D**) Representative Western blots to show expression of wild type R54L (lane 1) and R54L-4A (puromycin-resistant cell population; lane 3) in HeLa R54L KO cells. Loading control: ⍺-Tubulin. (**E**) ClustalW sequence alignment of the N-terminal domains of R54L from human (Homo sapiens), chimpanzee (Pan troglodytes), mouse (Mus musculus), and bovine (Bos taurus). Basic residues are shown in blue, and acidic residues are shown in red. The pink/grey box indicates basic residues mutated to alanines, and the green box indicates CDK2 phosphorylation consensus sequence and mutated serine to glutamate, as previously described (29). (**F**) Dot plot with medians of IdU tract lengths in HU-treated HeLa R54L KO cells expressing wild type R54L (WT) or R54L BM mutants (n=3; at least 100 fiber tracts/experiment analyzed). (**G**) Schematic of the DNA fiber assay protocol used in H. (**H**) Dot plot with medians of IdU tract lengths in CPT-treated HeLa or R54L KO cells expressing R54L BM mutants (n=3; at least 100 fiber tracts/experiment analyzed). IdU tract lengths were analyzed by Kruskal-Wallis test followed by Dunn’s multiple comparisons test (ns, not significant; *, p<0.05; ****, p<0.0001). R54L: RAD54L.

### RAD54L’s branch migration attribute contributes to its engagement in fork restraint

Fork reversal is a two-step process, in which fork regression is followed by BM to drive extensive reversal (60). RAD54L catalyzes BM, and its N-terminal domain is essential for this activity (29). As N-terminal mutations in RAD54L that selectively inhibit the BM activity have been described (29), we asked if RAD54L BM mutants would show defects in fork restraint. To this end, we expressed mutant RAD54L-S49E (deficient in oligomerization (29)), RAD54L-4A (deficient in binding to HJ-like DNA structures (29)), and RAD54L-4A/S49E (deficient in both binding to HJs and oligomerization (29)) in RAD54L KO cells (Fig. 5C-E). To assess defects in fork restraint, we chose the protocol as depicted in Fig. 5A and compared the response of RAD54L KO cells expressing mutant RAD54L to that of RAD54L KO cells with and without wild type RAD54L. As shown above (Fig. 2E, G) and reproduced herein independently, RAD54L KO cells exhibit a significant defect in fork restraint in the presence of low concentrations of HU (p<0.001; Fig. 5F; Fig. S5E). Fork restraint also is significantly impaired in RAD54L KO cells expressing RAD54L-S49E, RAD54L-4A, or the compound mutant RAD54L-4A/S49E (Fig. 5F; Fig. S5E), demonstrating that RAD54L BM activity is required for fork restraint. As expected, RAD54L ATPase activity is required for fork restraint, as IdU tract lengths in RAD54L KO cells expressing the ATPase dead mutant RAD54L (K189R) (61) are not significantly different from those in RAD54L KO cells (p=0.343 and p>0.999 for GRT #1 and GRT#2, respectively; Fig. S5F-I).

Next, we wondered if the defect in fork restraint of the RAD54L BM mutants could also be observed after treatment of cells with another replication stress-inducing drug. To do so, we treated cells with sublethal concentrations of camptothecin (CPT; Fig. 5G), a topoisomerase 1 inhibitor, shown to induce fork slowing and reversal, and- under the conditions tested here- no DSB formation (62). In CPT, RAD54L-4A and -4A/S49E expressing RAD54L KO cells showed a significant defect in fork restraint (p<0.0001) that was not significantly different from that of the RAD54L KO cells (p=0.437; Fig. 5H; Fig. S5J). Together, these results lead us to conclude that RAD54L BM activity is critical for fork restraint under conditions of mild replication stress.

### The requirement in fork restraint for RAD54L branch migration is specific to the FBH1 pathway

To dissect if RAD54L’s BM activity is required in both the FBH1 and the HLTF/SMARCAL pathways, we analyzed IdU tract lengths in the presence of 25 µM HU (Fig. 5A). RAD54L KO cells and RAD54L KO cells expressing wild type RAD54L or RAD54L-4A/S49E and with or without FBH1 or HLTF knockdown were subjected to mild replication stress (Fig. 6A). In accord with our earlier observations (Fig. 5B), IdU tract lengths were similar in all cell types with HLTF knockdown (Fig. 6B; Fig. S6A), suggesting that loss of fork reversal in the absence of HLTF is independent of RAD54L BM activity. However, after FBH1 knockdown, IdU tract lengths in RAD54L KO cells and RAD54L KO cells expressing the RAD54L-4A/S49E mutant were similar (p>0.999) but significantly shorter than those after FBH1 knockdown in RAD54L KO cells with wild type RAD54L (p<0.0001). Together, this suggests that RAD54L BM activity is essential in the FBH1 pathway, and the consequences of FBH1 loss in RAD54L-4A/S49E BM-deficient cells mirror those of FBH1 loss in RAD54L KO cells.

**Fig. 6.**
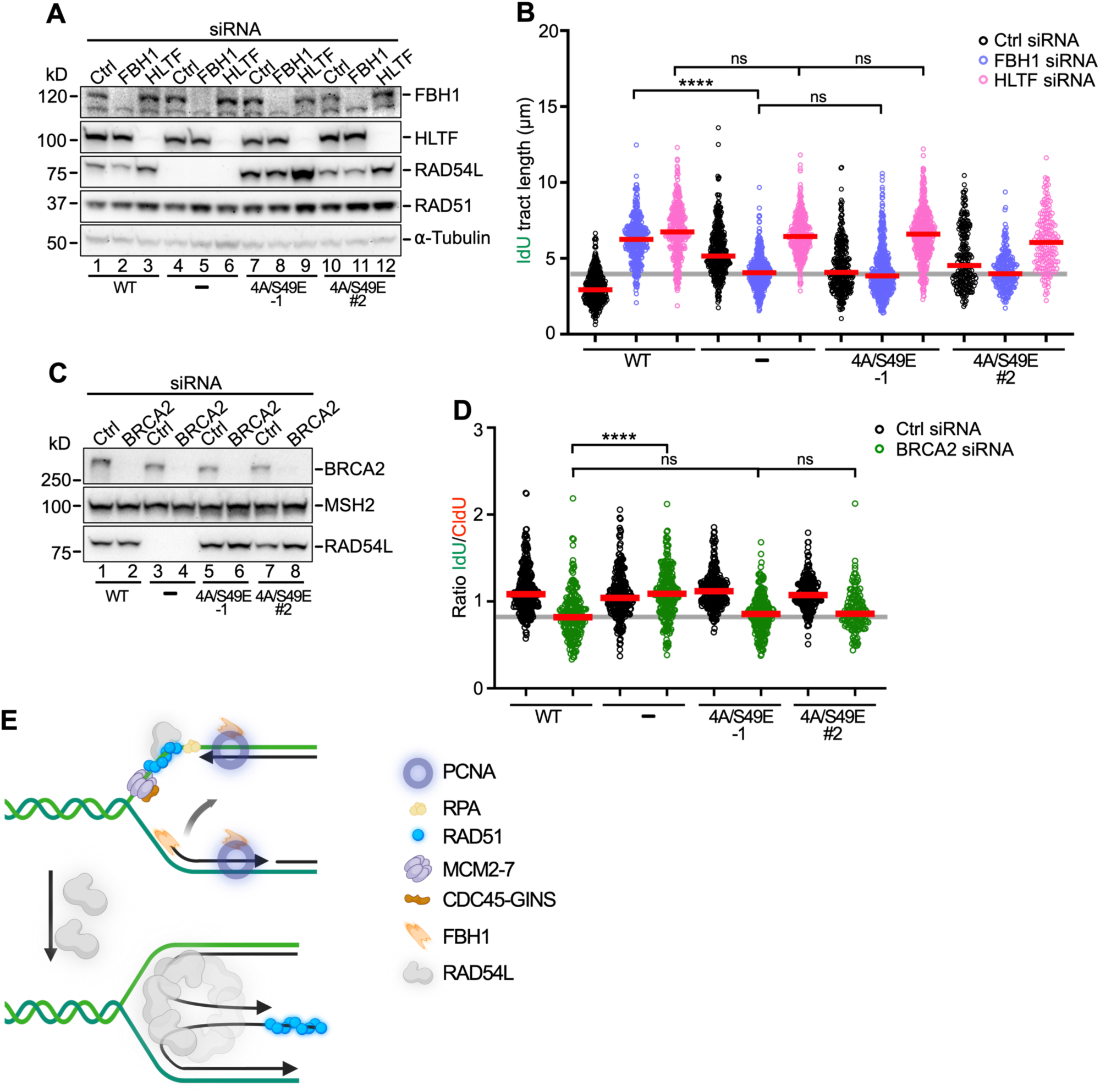
RAD54L’s branch migration activity is specifically required for the FBH1 pathway of RAD51-mediated fork reversal. (**A**) Representative Western blots to show extent of FBH1 and HLTF knockdown in R54L KO cells (lanes 4-6) and R54L KO cells expressing wild type RAD54L (lanes 1-3) or mutant R54L-4A/S49E (lanes 7-12). R54L-4A/S49E-1 dignifies a clonal isolate; R54L-4A/S49E #2 dignifies a puromycin-resistant cell population. Loading control: ⍺-Tubulin. (**B**) Dot plot with medians of IdU tract lengths in HU-treated (see Fig. 5A) R54L KO cells and R54L KO cells expressing wild type R54L (WT) or R54L-4A/S49E and transfected with Ctrl, FBH1, or HLTF siRNA (n=3; at least 100 fiber tracts/experiment analyzed; n=1 for RAD54L-4A/S49E #2). (**C**) Representative Western blots to show extent of BRCA2 knockdown in R54L KO cells (lanes 3-4) and R54L KO cells expressing wild type R54L (lanes 1-2) or mutant R54L-4A/S49E (lanes 5-8). R54L-4A/S49E-1 dignifies a clonal isolate; R54L-4A/S49E #2 dignifies a puromycin-resistant cell population. Loading control: MSH2. (**D**) Dot plot with medians of IdU tract lengths in HU-treated (see Fig. 5A) R54L KO cells and R54L KO cells expressing wild type R54L (WT) or R54L-4A/S49E and transfected with Ctrl or BRCA2 siRNA (n=3; at least 100 fiber tracts/experiment analyzed). (**E**) Model to explain the role of the RAD54L translocase in the FBH1 pathway of RAD51-mediated fork reversal. RAD54L may be recruited through its interaction with RAD51 on extended parental ssDNA. FBH1 is recruited to stalled replication forks by PCNA (74). Unwinding of lagging strand DNA through FBH1 (71) may initiate nascent strand annealing. RAD54L may then oligomerize on a 4-way junction and drive reversal through BM. The dependency of FBH1 on RAD54L may be indicative of coordinated recruitment and/or post-translational modification orchestrating sequential coaction. The exact number of subunits present in the RAD54L oligomer remain to be determined. Schematic created with Biorender.com.

We then tested RAD54L KO cells with wild type RAD54L or RAD54L-4A/S49E in the fork attrition assay (Fig. 4A) and with or without BRCA2 knockdown (Fig. 6C). In accord with our earlier observations (Fig. 4C), IdU/CldU ratios were >1 (indicative of fork protection) in RAD54L KO cells with BRCA2 knockdown (p<0.0001; Fig. 6D; Fig. S6B). In contrast, IdU/CldU ratios in RAD54L KO cells with RAD54L-4A/S49E and BRCA2 knockdown were comparable to those in cells with wild type RAD54L (p>0.999) and significantly lower than in RAD54L KO cells with BRCA2 knockdown (p<0.0001). These results show that the RAD54L BM mutant behaves like wild type RAD54L in the HLTF-mediated fork reversal pathway, in which the BRCA2 protein was shown to function as a major fork protection factor (10). These results suggest that loss of a RAD54L function other than its BM activity is required to prevent fork attrition in the absence of BRCA2 in the HLTF pathway.

## DISCUSSION

Our work establishes that RAD54L activity decelerates replication fork progression in human cells. Our results corroborate biochemical evidence (28) and are in strong support of RAD54L’s engagement in replication fork reversal in human cells. In biochemical assays (28), purified RAD54L was shown to use its BM activity to catalyze both the regression and restoration of model replication forks. In the presence of RAD51, however, as would be the situation in cells, the reaction was shown to be shifted toward the accumulation of the chicken foot structure (28). Our results in cells are in accord with this *in vitro* investigation and now have added one more attribute to the multi-functional RAD54L protein-limiting replication stress through fork restraint in human cells.

We have dissected that RAD54L functions with disparate requirements in the two distinct (14) RAD51-mediated fork reversal pathways. We show that RAD54L’s ability to catalyze BM is critical for fork restraint when engaged in the FBH1 pathway. In the HLTF/SMARCAL1 pathway, however, RAD54L BM activity is dispensable, suggesting that RAD54L attributes other than its BM activity contribute to fork reversal here.

RAD54L BM activity relies on its ability to function as an ATPase and form high-order RAD54L oligomers on HJ-like structures (63,64). Importantly, the 4A, S49E, and S49E/4A mutants tested here interfere with RAD54L oligomerization and compromise BM activity but show no defect in ATP hydrolysis or in their ability to stimulate the RAD51-mediated strand exchange reaction (29). Consequently, we infer that the stimulation of strand invasion contributes to the engagement of RAD54L in the HLTF pathway, as suggested recently (9). This scenario could explain why fork degradation transpires in BRCA2-depleted RAD54L KO cells expressing RAD54L-4A/S49E in contrast to cells with BRCA2 knockdown and complete loss of RAD54L. Accordingly, under mild replication stress and with HLTF/SMARCAL1 knockdown, RAD54L status has no or only a marginal effect on IdU tract lengths. This would be expected if HLTF/SMARCAL1 functions prior to and largely independently of RAD54L. Nonetheless, RAD54L is required during later stages in the HLTF/SMARCAL1-mediated fork reversal reaction, as shown here and previously (9), and we have established that this RAD54L engagement is not related to its ability to drive BM.

Genetic and biochemical studies have delineated that fork reversal is catalyzed by three replication fork remodelers of the SNF2 family (SMARCAL1, ZRANB3, and HLTF) or by the FBH1 helicase (14,54,65,66). Loss of either SMARCAL1, ZRANB3, or HLTF was shown to block nascent strand degradation in BRCA1/2-deficent cells, results that suggested their cooperation and non-redundant activities in fork reversal (10). Further, current models posit that fork reversal is a dynamic reaction that involves orchestrated actions of several proteins with specialized substrate preference (10,66,67). For example, in biochemical reconstitution assays ZRANB3 and HLTF were shown to be highly efficient in BM activity, while SMARCAL1 was little efficient in this regard. Moreover, SMARCAL1 and ZRANB3 were comparably efficient in annealing RPA-coated DNA, while HLTF had much reduced capacity in this assay (66).

To date, much less is known with respect to the molecular details of the FBH1 pathway of RAD51-mediated fork reversal. However, it is known that FBH1’s functional helicase domain is needed for the catalysis of fork reversal, but its ubiquitin ligase activity is not (14,54). While BRCA1/2, FANCD2 (10,13,15,16) and other proteins (68–70) protect reversed forks generated by SMARCAL1, HLTF, and ZRANB3, reversed forks generated by FBH1 are protected by 53BP1, and loss of FBH1 rescues fork degradation in the absence of 53BP1 (14). Importantly, we show here that loss of RAD54L also rescues fork degradation in the absence of 53BP1. Moreover, under conditions of mild replication stress, the block of fork restraint in cells with FBH1 knockdown is significantly ameliorated in the absence of RAD54L, suggestive of a concerted action between FBH1 and RAD54L in fork reversal. In contrast, RAD54L does not affect fork restraint in cells depleted for SMARCAL1 or HLTF, as their activities in fork reversal do not depend on RAD54L. Unlike in the SMARCAL1/HLTF pathway, in which RAD54L-4A/S49E behaves identical to wild type RAD54L, RAD54L KO cells expressing RAD54L-4A/S49E mirror RAD54L KO cells in the FBH1 pathway. These results are in strong support of FBH1 relying on RAD54L to drive BM to catalyze fork reversal. We suggest a model in which the combined activities of RAD54L and FBH1 lead to fork reversal (Fig. 6E). In this model, FBH1 may help unwind the growing lagging strand to promote nascent strand annealing, as suggested previously (71), and RAD54L may regress the 4-way junction through its BM activity, as suggested by the results described here.

A growing body of work has provided evidence that replication-associated ssDNA gaps affect the response of cancer cells to chemotherapy, suggesting that enzymes that limit ssDNA gap accumulation may represent useful targets in cancer therapy (56,72,73). We propose that RAD54L may be one such target, as ssDNA gap formation is enhanced in its absence and further exacerbated through treatment of RAD54L-deficient cells with PARPi. This proposal may be particularly attractive, as RAD54L is also a key player in the HR DNA repair pathway, a pathway frequently associated with resistance to cancer therapy.

## Supporting information

Supplemental Tabels

## DATA AVAILABILITY

### SUPPLEMENTARY DATA

Supplementary Data are available at NAR online.

### AUTHOR CONTRIBUTIONS

Mollie E. Uhrig: Investigation, methodology, validation, formal analysis, writing of the manuscript. Neelam Sharma: Investigation, methodology, validation. Petey Maxwell: Validation. Platon Selemenakis: Critical reagents. Alexander V. Mazin: Conceptualization, funding acquisition. Claudia Wiese: Conceptualization, formal analysis, writing of the manuscript, funding acquisition.

### FUNDING

This work was supported by National Institutes of Health [R01GM144579, R01 GM136717, R01 CA237286], the Cancer Prevention and Research Institute of Texas (CPRIT) REI RR210023, and by the Congressionally Directed Medical Research Programs BC191160. A.V.M. is the holder of the Joe R. and Teresa Lozano Long Chair in Cancer Research. Funding for open access charge [NIH/R01GM144579].

