## Supplemental Material for "Disparate requirements for RAD54L in replication fork reversal"

**Supplementary Figure S1**
**Supplementary Figure S2**
**Supplementary Figure S3**
**Supplementary Figure S4**
**Supplementary Figure S5**
**Supplementary Figure S6**

**Supplementary Table S1**
**Supplementary Table S2**
**Supplementary Table S3**
**Supplementary Table S4**
**Supplementary Table S5**

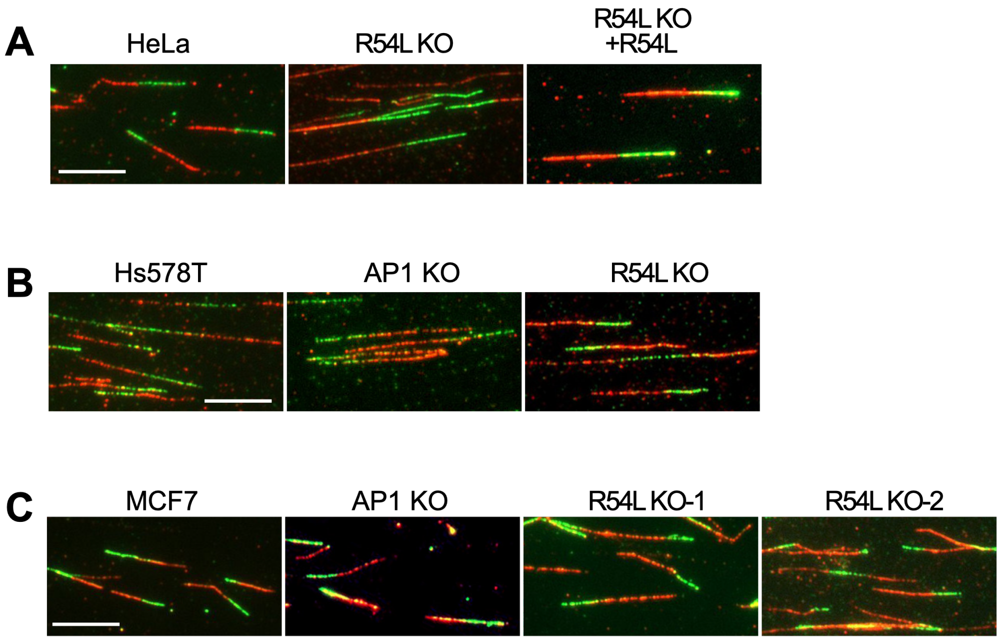


**Fig. S1** Related to Fig.1. **RAD54L restrains fork progression.** Representative micrographs of DNA fibers in unperturbed cell lines. (**A**) HeLa cells and derivatives. (**B**) Hs578T cells and derivatives. (**C**) MCF7 cells and derivatives. R54L: RAD54L; AP1: RAD51AP1. Scale bars: 10 µm.

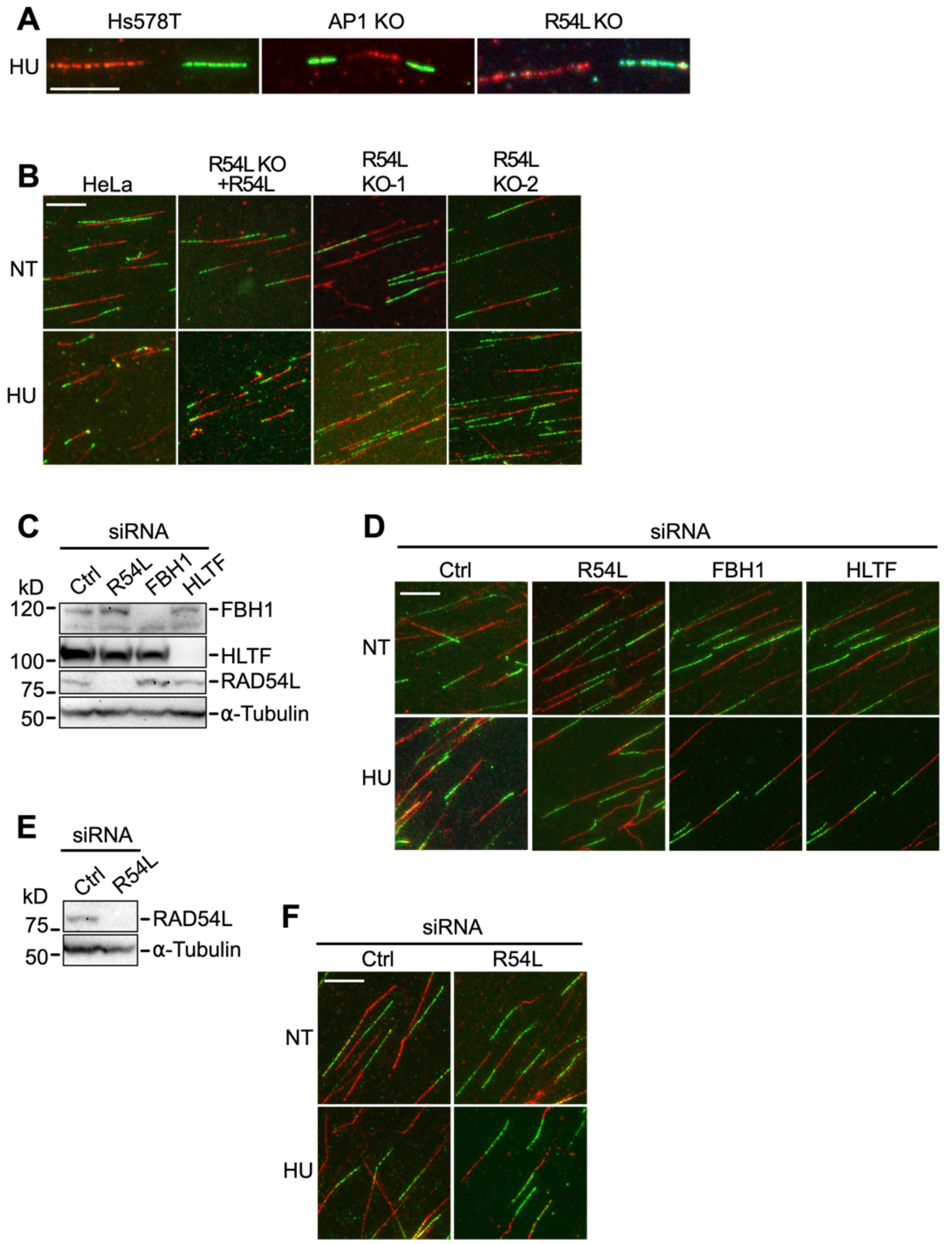


**Fig. S2** Related to Fig.2. **Loss of RAD54L does not affect replication restart and leads to longer replication tracts during mild replication stress.** (**A**) Representative micrographs of DNA fibers in Hs578T cell lines to determine replication restart by IdU tract lengths (green). (**B**) Representative micrographs of DNA fibers in HeLa cells and derivatives to determine the response of cells to mild replication stress (25 µM HU) by IdU tract lengths. NT: not treated. (**C**) Western blots to show the extent of protein knockdown in HeLa cells after transfection with Ctrl, R54L, FBH1 or HLTF siRNA. Loading control: α-Tubulin. (**D**) Representative micrographs of DNA fibers in HeLa cells transfected with siRNA as indicated, to determine the response of cells to mild replication stress by IdU tract lengths. NT: not treated. (**E**) Western blots to show the extent of R54L knockdown in hTERT RPE-1 after transfection with Ctrl or R54L siRNA. Loading control: α-Tubulin. (**F**) Representative micrographs of DNA fibers in hTERT RPE-1 cells transfected with Ctrl or R54L siRNA to determine the response of cells to mild replication stress by IdU tract lengths. NT: not treated. R54L: RAD54L; AP1: RAD51AP1. Scale bars: 10 µm.

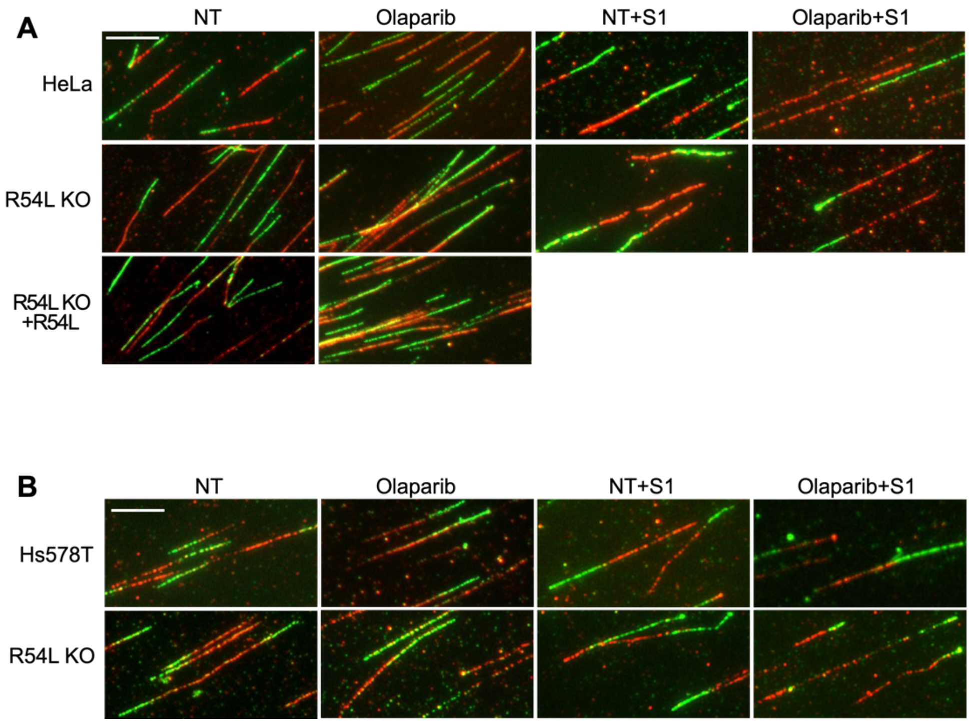


**Fig. S3** Related to Fig.3. **Loss of RAD54L accelerates replication fork progression and ssDNA gap formation.** (**A**) Representative micrographs of DNA fibers in HeLa cells and derivatives without (NT) and after a 24h-treatment with Olaparib (10 µM) with and without S1 nuclease digest. (**B**) Representative micrographs of DNA fibers in Hs578T cell lines without (NT) and after a 24h-treatment with Olaparib (10 µM) with and without S1 nuclease digest. R54L: RAD54L. Scale bars: 10 µm.

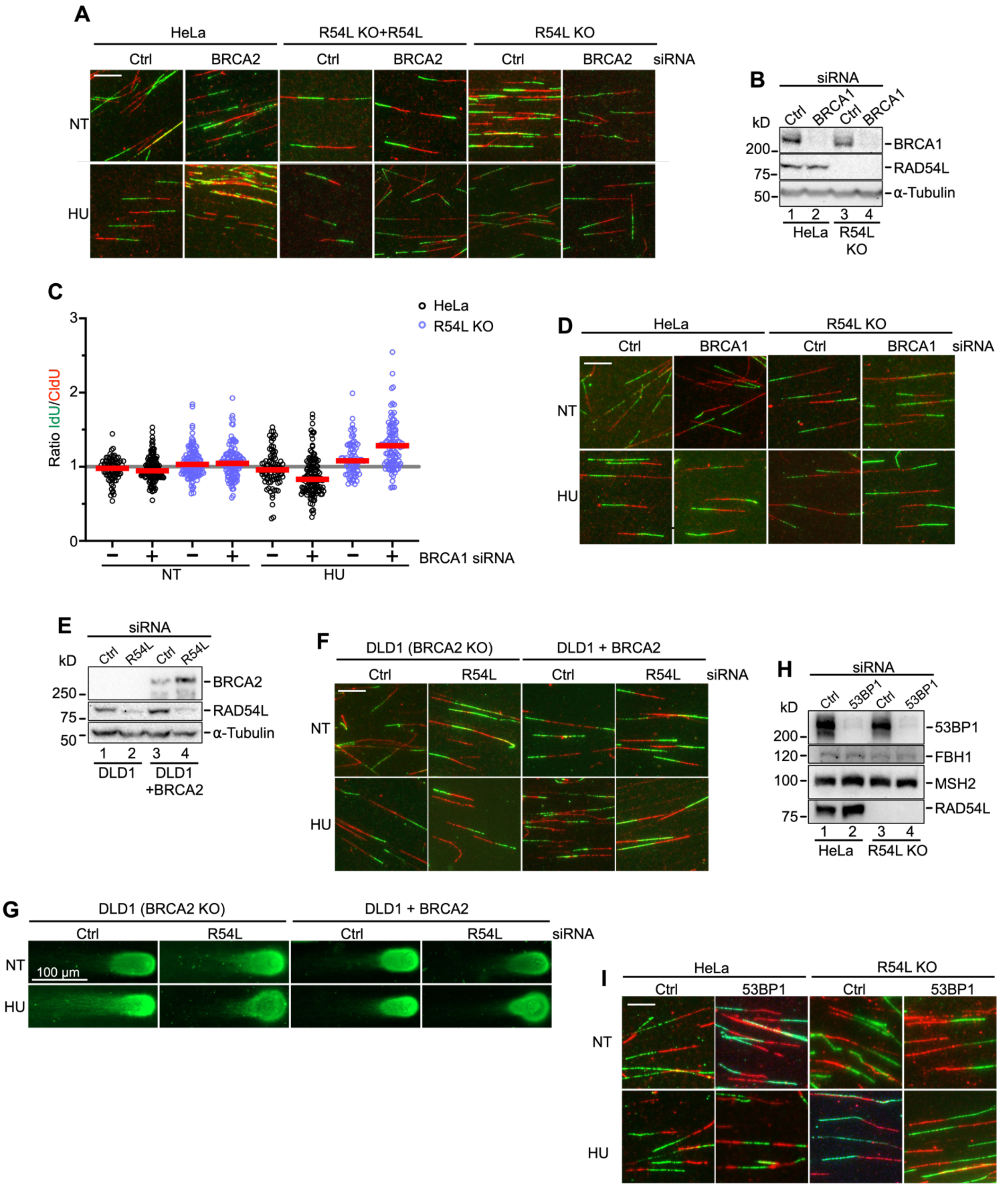


**Fig. S4** Related to Fig. 4. **Loss of RAD54L prevents fork degradation in both BRCA2- and 53BP1-deficient cells.** (**A**) Representative micrographs of DNA fibers in HeLa cells and derivatives transfected with Ctrl or BRCA2 siRNA. (**B**) Representative Western blots to show extent of BRCA1 knockdown in HeLa and R54L KO cells for the experiment shown in C. Loading control: α-Tubulin. (**C**) Dot plot with medians of IdU/CldU tract length ratios in HeLa and R54L KO cells transfected with Ctrl (**-**) or BRCA1 siRNA (n=1; at least 100 fiber tracts/condition analyzed). (**D**) Representative micrographs of DNA fibers in HeLa and R54L KO cells transfected with Ctrl or BRCA1 siRNA. (**E**) Western blots to show extent of R54L knockdown in DLD1 and DLD1+BRCA2 cells. Loading control: α-Tubulin. (**F**) Representative micrographs of DNA fibers in DLD1 and DLD1+BRCA2 cells transfected with Ctrl or R54L siRNA. (**G**) Representative micrographs of DLD1 and DLD1+BRCA2 cells transfected with Ctrl or R54L siRNA with eluted DNA after neutral comet assay. (**H**) Western blots to show extent of 53BP1 knockdown in HeLa and R54L KO cells. Loading control: α-Tubulin. (**I**) Representative micrographs of DNA fibers in HeLa and R54L KO transfected with Ctrl or 53BP1 siRNA. R54L: RAD54L. NT: not treated. Scale bars: 10 μm.

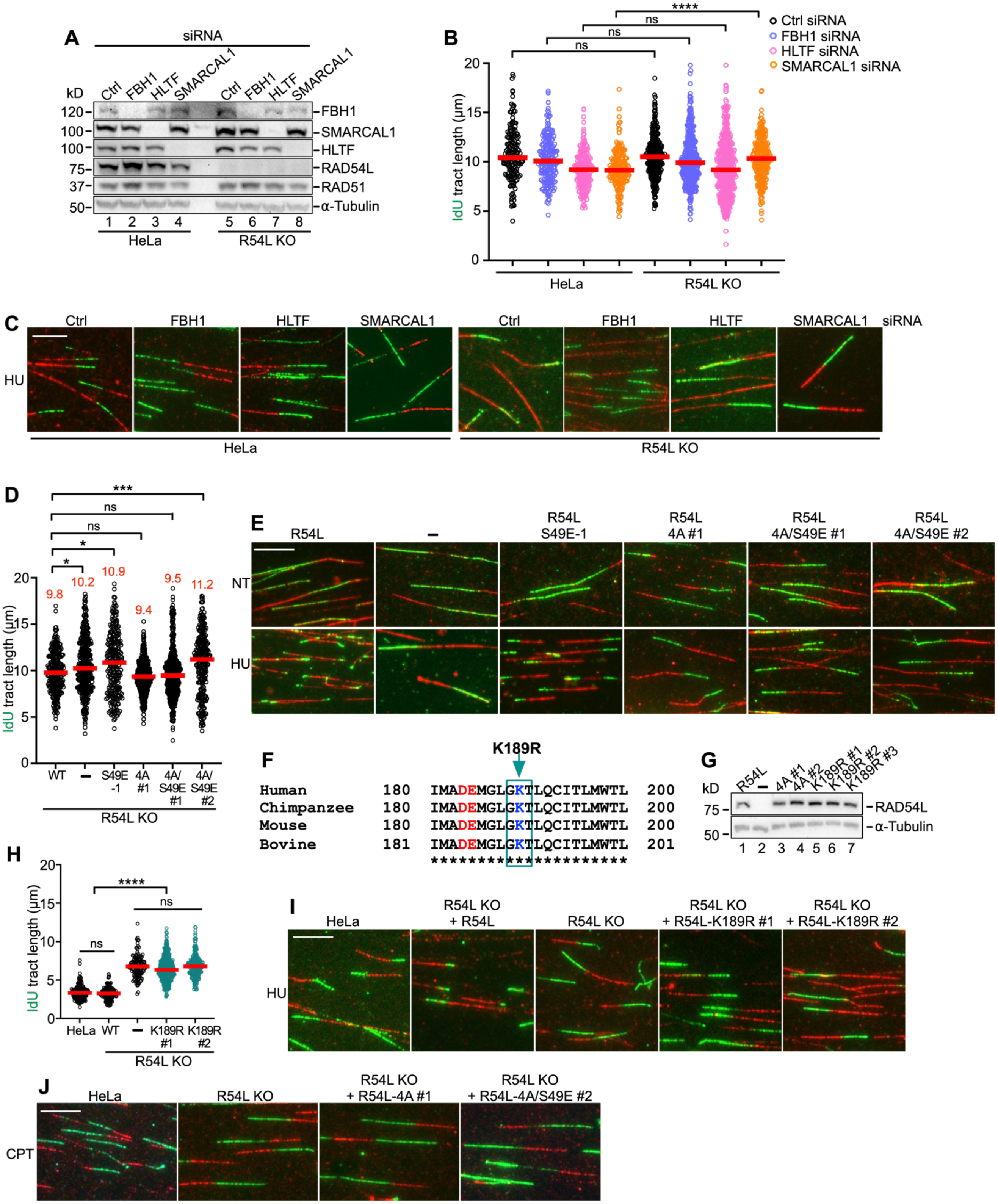

**Fig. S5** Related to Fig. 5. **RAD54L's branch migration activity contributes to its ability to restrain fork progression.** (**A**) Western blots to show the extent of protein knockdown after transfection of HeLa and R54L KO cells with Ctrl, FBH1, HLTF, or SMARCAL1 siRNA. Loading control: α-Tubulin. (**B**) Dot plot with medians of IdU tract lengths in unperturbed HeLa and R54L KO cells transfected with siRNAs, as indicated. (**C**) Representative micrographs of DNA fibers in HeLa and R54L KO cells transfected with siRNAs as indicated, and exposed or not (NT) to HU. (**D**) Dot plot with medians of IdU tract lengths in unperturbed R54L KO cells (-) and R54L KO cells expressing wild type R54L (WT) or RAD54L BM mutants. R54L-S49E-1 is a clonal isolate; R54L-4A #1 and R54L-4A/S49E #1/#2 expressing cells are puromycin-resistant cell populations. (**E**) Representative micrographs of R54L KO cells and R54L KO cells expressing R54L wildtype or BM mutants and treated or not (NT) with HU. (**F**) ClustalW sequence alignment of the GKT box encompassing region of RAD54L from human (Homo sapiens), chimpanzee (Pan troglodytes), mouse (Mus musculus), and bovine (Bos taurus). Basic residues are shown in blue, and acidic residues are shown in red. The green box indicates the GKT box and mutated lysine to arginine. (**G**) Western blots to show the expression levels of wild type R54L (clonal isolate), R54L-4A, and R54L-K189R protein (independently isolated mixed cell populations) in R54L KO (**-**) cells. Loading control: α-Tubulin. (**H**) Dot plot with medians of IdU tract lengths in HU-treated R54L KO (**-**) cells and R54L KO cells expressing wild type R54L (WT) or mutant R54L-K189R. (**I**) Representative micrographs of DNA fibers for the experiment shown in H. (**J**) Representative micrographs of DNA fibers for HeLa, R54L KO cells, and R54L KO cells expressing R54L BM mutants exposed to CPT (25 nM). IdU tract lengths were analyzed by Kruskal-Wallis test followed by Dunn's multiple comparisons test (ns, not significant; ∗, p<0.05; ∗∗∗, p<0.001; ∗∗∗∗, p<0.0001). R54L: RAD54L. Scale bars: 10 µm.

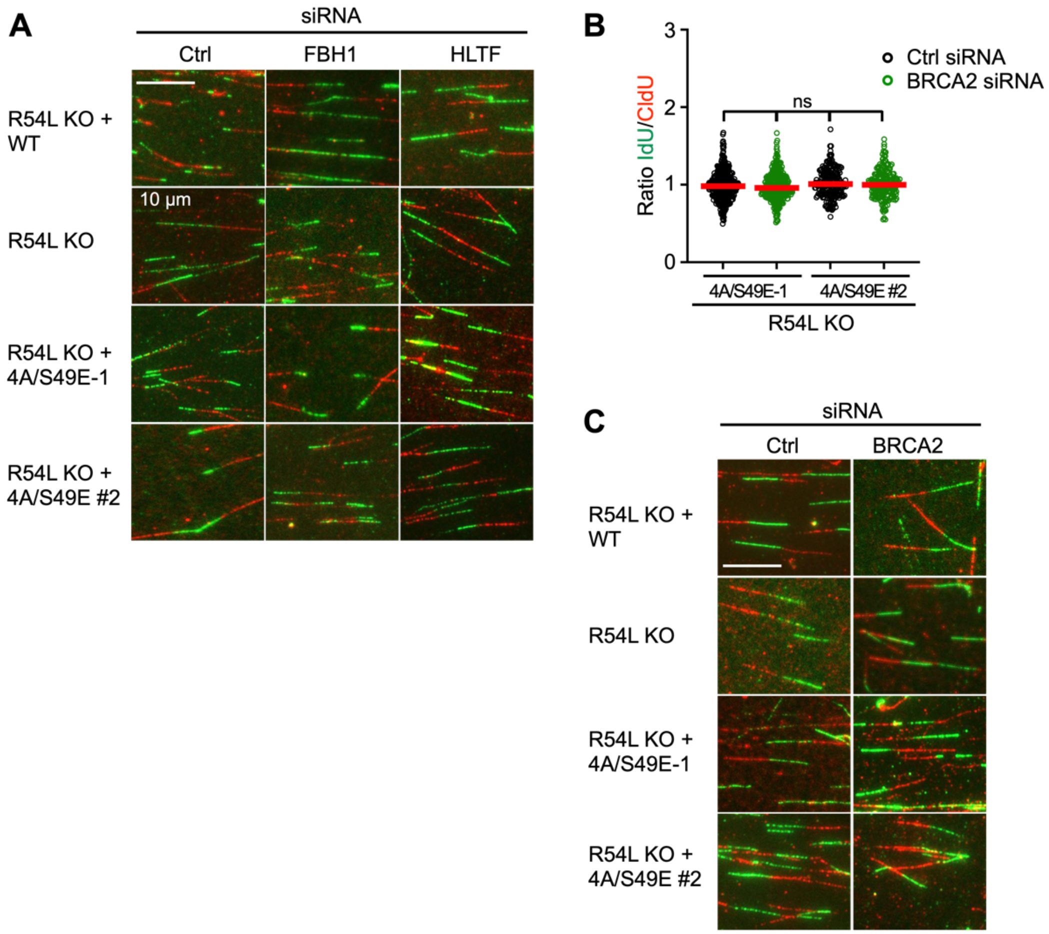


**Fig. S6** Related to Fig. 6. **RAD54L's branch migration activity is specifically required for the FBH1 pathway of RAD51-mediated fork reversal.** (**A**) Representative micrographs of R54L KO cells and R54L KO cells expressing R54L wildtype or mutant R54L defective in BM transfected with Ctrl, FBH1, or HLTF siRNA and treated with HU. (**B**) Dot plot with medians of IdU/CldU tract length ratios in unperturbed R54L KO cells expressing mutant R54L-4A/S49E transfected with Ctrl or BRCA2 siRNA. R54L-4A/S49E-1 dignifies a clonal isolate; R54L-4A/S49E #2 dignifies a puromycin-resistant cell population. (**C**) Representative micrographs of R54L KO cells and R54L KO cells expressing R54L wildtype or mutant R54L defective in BM transfected with Ctrl, FBH1 or BRCA2 siRNA and treated with HU. R54L: RAD54L. Scale bars: 10 µm.
