## Supplemental Tabels for "Disparate requirements for RAD54L in replication fork reversal"

Supplementary Table S1. siRNAs used in this study.

| **Name** | **Target Sequence (listed 5´- 3´)** | **Reference** |
| --- | --- | --- |
| non-depleting control (Ctrl) | GATTCGAACGTGTCACGTCAA | [1] |
| RAD54L | AAGCATTTATTCGAAGCATTT | [2] |
| BRCA1 | AAGCTCCTCTCACTCTTCAGT | [3] |
| BRCA2 | TTGGAGGAATATCGTAGGTAA | [4] |
| HLTF | Pool of 3 target-specific siRNAs (Santa Cruz Biotechnology) | sc-45943 |
| FBH1 | Pool of 3 target-specific siRNAs (Santa Cruz Biotechnology) | sc-90469 |

Supplementary Table S2. sgRNAs used in this study.

| **Name** | **Target** | **Sequence (listed 5´- 3´)** |
| --- | --- | --- |
| sgRNA A | *RAD54L* (exon 8) | CGGAGGGAGGATCCAACCTC |
| sgRNA B | *RAD54L* (exon 8) | GTACCAGTTCTTCACCAGGC |
| sgRNA (1) | *RAD51AP1* (exon 2*)* | TTTGACCACTCTGACAGTGA |
| sgRNA (2) | *RAD51AP1* (exon 3*)* | AACCTAACTTGAACAATCTC |
| sgRNA (3) | *RAD51AP1* (exon 5/6*)* | AGTGTAGCCAGTGATTATTT |

Supplementary Table S3. PCR primers used in this study for amplification of genomic DNA.

| **Primer** | **Target** | **Sequence (listed 5´- 3´)** | **Product Length (bp)** |
| --- | --- | --- | --- |
| P1 | *RAD54L* (intron 7) | AGACTACCATCCCTGGGACA | 597 |
| P2 | *RAD54L* (intron 8) | CAACAGAAAAGGTGTAAAGGGAACA |  |
| P1 | *RAD51AP1* (intron 1) | TCCCCGCGGTAAAATGCAAATC | 496 |
| P2 | *RAD51AP1* (intron 2) | CACCTGGCCTGTTCATTTATCACC |  |
| P3 | *RAD51AP1* (intron 2) | AGGGCACAAAAACAAAAGTCGA | 470 |
| P4 | *RAD51AP1* (intron 3) | CGTCCTGTTTTCTGACTGCACC |  |
| P5 | *RAD51AP1* (intron 4/5) | TGCCAGTTGGAGTTTGGGATCA | 420 |
| P6 | *RAD51AP1* (intron 5/6) | AAGCCACGGGTAGTTATGACCC |  |

Supplementary Table S4. Oligonucleotides used in this study for site-directed-mutagenesis.

| **Template** | **Oligonucleotide** | **Target Sequence (listed 5´- 3´)** |
| --- | --- | --- |
| pENTR1A-RAD54L [5] | S49E forward | GTGTTTCCTGgagCCTTTTCGGAAACC |
|  | S49E reverse | TCCTGGATCTGGGTCTCA |
| pENTR1A-RAD54L-S49E | 33AA34 forward | AGTGACTCCTgctgcaCGGAAATCCAGCAGTG |
|  | 33AA34 reverse | AGGCCAGGTTGCCAGTCT |
| pENTR1A-RAD54L-33AA34-S49E | 35AA36 forward | TCCTGCTGCAgctgcaTCCAGCAGTGAGACC |
|  | 35AA36 reverse | GTCACTAGGCCAGGTTGC |

Mutated bases in lower case characters.

**Supplementary Table S5.** Sequencing results for genotyping of *RAD54L* and *RAD51AP1* disrupted MCF7 cells.

| **Gene, estimated copy number*****, chromosome** | **Cell line** | **Genomic sequence (gRNA-PAM)^#^** | **Indel (nt)** |
| --- | --- | --- | --- |
| *RAD54L*  3  Chr. 1 | R54L KO-1 | GCCTT**CCAGCCTGGTGAAGAACTGGTAC**AATGAGGTTGGGAAATGGCT**CGGAGGGAGGATCCAACCTCTGG**CCATC | WT |
|  |  | GCCTT**CCAGCCTGGTGAAGAACTGGTAC**AATGAGGTTGGGAAATG**----------------CAACCTCTGG**CCATC | -16 |
|  |  | GCCTT**CCAGCCTGGT-----------AC**AATGAGGTTGGGAAATG----------------**CAACCTCTGG**CCATC | -27 |
|  |  | GCCTT**CC-----------------------------------------------------CCAACCTCTGG**CCATC | -53 |
| *RAD54L*  3  Chr. 1 | R54L KO-16 | GCCTT**CCAGCCTGGTGAAGAACTGGTAC**AATGAGGTTGGGAAATGGCT**CGGAGGGAGGATCCAACCTCTGG**CCATC | WT |
|  |  | GCCTT**CCAGCCTGGTGAAGAACTGGTAC**AATGAGGTTGGGAAATG**----------------CAACCTCTGG**CCATC | -16 |
|  |  | GCCTT**CCAGCC--------------------------------------------------------------**ATC | -62 |
|  |  | GCCTT**CCAGCCTGGT-----------AC**AATGAGGTTGGGAAATG**----------------CAACCTCTGG**CCATC | -27 |
| *RAD51AP1*  3  Chr. 12 | AP1 KO-22 | TCATAGACATAAGAAACCAGTCAATTACTCACAG**TTTGACCACTCTGACAGTGATGG**TAAGTAAAGCTTCTTATTT | WTE2 |
|  |  | TCATAGACATAAGAAACCAGTCAATTACTCACAG**TTTGACCACTCTGACAG**ag**TGATGG**TAAGTAAAGCTTCTTATTT | +2 |
|  |  | GGAGTTAAAACAAGATAAACCAA**AACCTAACTTGAACAATCTCCGG**AAAGAAGAAATCCCAGTACAAGAGAAAACC | WTE3 |
|  |  | GGAGTTAAAACAAGATAAACCAA**AACCTAACTTGAACAAT-**a**CCGG**AAAGAAGAAATCCCAGTACAAGAGAAAACC | -1 |
|  |  | AAGTCTCCTCATATCTCTAATTGC**AGTGTAGCCAGTGATTATTTAGG**TAAGTTTTTTATATTAATAATTTTTCTCA | WTE6 |
|  |  | AAGTCTCCTCATATCTCTAATTGC**AGTGTAGCCAGTGATTATTT**t**AGG**TAAGTTTTTTATATTAATAATTTTTCTCA | +1 |

*Estimated from: [6] .

**^#^**Small lettering for insertions and bp changes.

WT: wild-type sequence.
